# ModelSEED v2: High-throughput genome-scale metabolic model reconstruction with enhanced energy biosynthesis pathway prediction

**DOI:** 10.1101/2023.10.04.556561

**Authors:** José P. Faria, Filipe Liu, Janaka N. Edirisinghe, Nidhi Gupta, Samuel M.D. Seaver, Andrew P. Freiburger, Vibhav Setlur, Qizhi Zhang, Pamela Weisenhorn, Neal Conrad, Raphy Zarecki, Hyun-Seob Song, Matthew DeJongh, Aaron A. Best, Robert W. Cottingham, Adam P. Arkin, Christopher S. Henry

**Affiliations:** Data Science and Learning Division, Argonne National Laboratory, Lemont, Illinois, USA; Biosciences Division, Argonne National Laboratory, Lemont, Illinois, USA; Mathematics and Computer Science Division, Argonne National Laboratory, Lemont, Illinois, USA; Newe Ya’ar Research Center, Agricultural Research Organization, Ramat Yishay, Israel; Departments of Biological Systems Engineering and Food Science and Technology, Nebraska Food for Health Center, University of Nebraska–Lincoln, Lincoln, Nebraska, USA; Department of Computer Science, Hope College, Holland, Michigan, USA; Department of Biology, Hope College, Holland, Michigan, USA; Biosciences Division, Oak Ridge National Laboratory, Oak Ridge, Tennessee, USA; Environmental Genomics and Systems Biology Division, E.O. Lawrence Berkeley National Laboratory, Berkeley, California, USA

**Author notes:** José P. Faria and Filipe Liu contributed equally to this work.

## Abstract

Since the release of ModelSEED in 2010, the systems biology research community has used the ModelSEED genome-scale metabolic model reconstruction pipeline to build over 200,000 draft metabolic reconstructions that support hundreds of publications. Here we describe the first comprehensive update to this reconstruction tool, with new features such as (i) a dramatically improved representation of energy metabolism, which ensures that models produce accurate amounts of ATP per mol of nutrient consumed; (ii) a new template for Archaea model reconstruction; and (iii) a significantly improved curation of all metabolic pathways with mappings to RAST subsystems annotations. We applied the improved pipeline to build and analyze core and genome-scale models for Archaea and Bacteria genomes in KEGG. The new ModelSEED pipeline generates larger models that possess more reactions and genes and require fewer gap-filled reactions. In addition, we see conserved patterns in the ATP biosynthesis mechanism across phylogeny, and identify clades where our understanding of energy biosynthesis is still poor. ModelSEED v2 is available through KBase and the ModelSEED website (https://modelseed.org/), which supports reconstruction, gap-filling, flux balance analysis and model export through a modern web interface.

## Introduction

We are entering the fourth decade of constraint-based metabolic modeling research that was kick-started by the seminal studies of *Escherichia coli* by Varma et al. (1, 2). Genome-scale metabolic models (GEMs) have become a standard research tool in many areas of systems biology research (3, 4). GEM reconstruction was originally a manual process involving gathering of data, laborious refinement, and network evaluation (5). While manual curation and refinement of GEMs remain crucial, the release of the ModelSEED pipeline in 2010 ushered in a new era of automated GEM reconstruction (6), which enabled researchers to obtain a draft model quickly and therefore focus their efforts on model refinement and simulation analysis. Additional GEM reconstruction pipelines and toolboxes have become available over the past decade (7–11), which are contrasted in a recent systematic review (12). Another recent survey on the availability of GEMs documented reconstructions for over 6,000 organisms (4) including extensive model reconstruction efforts such as AGORA (13, 14) and CarveMe (9) and hundreds of curated models in resources such as BIGG (15) and BioModels (16). With the accessibility and ease of use of these new pipelines, researchers rapidly moved from modeling single organisms to modeling communities with multiple species (17, 18). These advances in metabolic reconstruction, concurrent with advances in metagenome sequencing, assembly and binning to produce metagenome-assembled genomes (MAGs), are laying the foundation for impactful discoveries in microbial ecology (19).

The development of MAGs creates new challenges and opportunities for GEM reconstruction and GEM-based analyses. For example, GEMs can help assess the quality of MAGs (20) and even help fill gaps through MAG-based function analyses, demonstrated by using the original ModelSEED to generate draft models for an extensive compendium of gut microbiome (13, 14) and environmental MAGs (20, 21). As the community debates the reliability of MAGs (22), modeling pipelines must adapt by becoming more proficient in modeling lower-quality and incomplete genomes. Increasing robustness requires new tools to accurately classify organisms before reconstruction and modularize pipelines that empower community contributions and updates to specific components as developments are made.

We therefore developed ModelSEED v2 (MS2) to tackle these new challenges and rapidly build accurate genome-scale models for ever-increasing numbers of genomes and MAGs. MS2 is accessible through KBase and through the redesigned ModelSEED website, whose open-source backend is available at https://github.com/ModelSEED/modelseed-api. We improved the ModelSEED reconstruction procedure by constructing core metabolism models of the central carbon metabolism and energy biosynthesis pathways (23), testing them for proper ATP production and ensuring that ATP production does not excessively increase when expanding the core model to a genome-scale model or during the gap-filling process. Although excellent approaches do exist to correct ATP overproduction in models during downstream analysis (24–26), these are plagued with uncertainty over which ATP mechanisms should be kept or discarded, and they disrupt only unlimited ATP production loops as opposed to overly high ATP production caused by erroneous gap-filling (e.g., an anaerobic organism growing aerobically). For this reason, we posit that it is essential to ensure correct ATP production mechanisms during the initial reconstruction process to ensure that ATP overproduction does not occur in the first place. Furthermore, to handle the necessary expansion of templates, we developed machine learning (ML) classifiers that automatically determine and apply the correct template to each new genome reconstruction (manual template selection prior to reconstruction is also available) (27). This ML approach and the ability to generate new classifiers permit the rapid introduction of additional modeling templates.

The accuracy of metabolic pathway reconstruction in the MS2 pipeline was improved by updating the ModelSEED biochemistry database (28) to include the latest reaction data from KEGG (29), MetaCyc (30), BIGG (15), and published models. Additionally, we manually curated the central pathways in our reconstruction templates to reconcile pathway representation across these multiple databases (23), and then curated our mapping of RAST (31) functional roles to this reconciled biochemistry based on data mined from KEGG and published metabolic models, further improving reconstruction accuracy. Within the KBase platform (32), we demonstrate the improved MS2 pipeline by constructing draft GEMs for Bacteria and Archaea genomes in KEGG. We show how the gene counts and modeling metrics (ATP production, biomass yields, reaction classification, pathway representation) improved with this new release of the ModelSEED. Moreover, we analyze energy biosynthesis for the entire collection of Archaea and Bacteria models and identify conservation with phylogeny. A qualitative and quantitative comparison study with the CarveMe (9) pipeline (a top-performing pipeline based on recent assessment (12)) for the same KEGG genomes revealed agreement in many circumstances as well as strengths and weaknesses of each pipeline that may be useful for specific contexts. The MS2 pipeline is available on KBase (kbase.us). Details on the KBase application, including a tutorial on how to use the MS2 App, are available in *Supplementary KBase Narrative 1*. ModelSEED is also available at https://modelseed.org/ through a redesigned web interface backed by an open-source REST API (https://github.com/ModelSEED/modelseed-api). The web platform supports model reconstruction, gap-filling, flux balance analysis, model editing, and export in ModelSEED JSON, SBML, and COBRA JSON formats.

## Methods

### Use of machine learning classifiers to identify specific model templates

We developed a set of machine learning models using the Python scikit-learn package (33) to classify prokaryotic genomes into specific microbial classes. We utilize these classes to automatically match the genome to our model reconstruction templates. Currently we support the following templates: Archaea, Bacteria gram-positive, and Bacteria gram-negative (27). These templates aid in deriving unique biochemistry and physiology for distinctive groups of organisms during the model construction process. Models are reconstructed with only one template based on the RAST annotations in the genome. Template order and hierarchy have no impact on this classification. The new ML (k-nearest neighbors) classifiers replace the naive Bayes classifiers used in the original ModelSEED for gram-positive and gram-negative Bacteria classification.

### Definition of template metabolic models

Model reconstruction templates contain all the data – biochemistry from the ModelSEED database (28), mappings of gene functional roles to reactions, and the biomass objective functions – that are required by the MS2 pipeline to build new metabolic models. The ModelSEED database comprehensively integrates other biochemical databases such as KEGG, MetaCyc, BIGG, and published models and includes many reactions that are not suitable or are mutually redundant for inclusion in metabolic models. Reactions in KEGG and MetaCyc have generic versions without electron donors and acceptors defined, which can obviate energy biosynthesis, and they often include a mixture of lumped and unlumped reactions, which can cause entire pathways to be bypassed during GEM simulation. Our MS2 model templates act as “filters” by including only “modeling-ready” reactions in models. Filtering criteria include (1) no abstract compounds such as “acceptor” and “donor” are allowed; (2) reaction must be mass and charged balanced; (3) all reactants must have defined charge and molecular formula (with few explicit exceptions such as biomass and APS); (4) highly lumped reactions should be avoided when unlumped alternatives exist; (5) reactants should be stereochemically explicit; and (6) all metabolites in the model should be standardized to their primary protonated form at pH 7.5. Other model reconstruction tools (9) do a similar “filtering” approach by using reactions only from the BIGG database, which is designed to include only reactions suitable for metabolic modeling. However, limiting models to only the BIGG database limits biochemical description to only a subset of organisms and, therefore, fails to capture the full diversity of microbial metabolism.

Equally important is the ability to use templates to curtail biochemistry by taxonomy. The original ModelSEED release contained only two templates – gram-positive and gram-negative – which ensured that gram-positive organisms would not be reconstructed with reactions or biomass components related to the gram-negative cell wall but constrained reconstruction accuracy to only Bacteria. MS2 introduces an Archaeal template and a core metabolism template that considerably broadens the reconstruction potential of our pipeline and further improves Bacteria gram-negative and gram-positive templates with new biochemical curation. A cyanobacteria-specific template is not currently available, and the results below demonstrate the resulting limitations for cyanobacterial reconstruction. The MS2 pipeline allows user-generated custom templates to address gaps in template coverage and curation. Custom templates can also introduce missing reactions, functional role mappings, and more specific biomass objective function definitions for a given phylogenetic group. The template repository is open-source; contributions are accepted via pull requests. These templates further limit gap-filling – the process of adding the fewest reactions to achieve model growth in a flux balance analysis (FBA) (34) simulation with a given media – to only those reactions that are in the template assigned to the reconstruction. This design prevents behavior where gap-filling could add Bacteria-specific reactions to fill gaps in a model for Archaea because the Bacterial pathway involves fewer reactions. Furthermore, template assignment ensures that as long as the organism is classified correctly, adequate reactions/pathways will be gap-filled. Organism classification is increasingly essential nowadays when extraction of MAGs is standard practice, given that the quality of such genomes can vary widely.

The aforementioned advantages are equally valid for separating templates into core and genome-scale templates. Core templates permit the isolation of energy biosynthesis pathways from the rest of the metabolism and ensure the proper functioning of electron transport chains (ETCs). However, reactions within these essential energy-generating processes can intertwine with other pathways at the genome-scale and consequently generate ATP production loops that create unrealistic model behavior. ATP production loops become even more problematic if gap-filling is applied since FBA can exploit these loops to achieve a mathematically optimal but biologically undesirable solution. For example, the reaction associated with the Archaea-specific version of the *Rnf* protein complex in acetoclastic methanogens (35) can couple with other energy biosynthesis reactions for infinite ATP production. We created taxa-specific templates to account for these edge cases when appropriate. The current core template ensures that MS2 produces “ATP-safe” models that lack these ATP production loops. Energy loops are prevented at the core level, and reactions are not added to a model if they lead to unrealistic ATP production during expansion to the genome-scale. The following sections provide more details on the reconstruction of core metabolism models and our “ATP-safe” reconstruction approach. The templates used for the analyses reported here are preserved in *Supplementary Data S1*. Versioned current and historical templates are also available from the ModelSEEDTemplates repository.

### Reconstruction of core metabolism metabolic models

Accurate representation of central carbon metabolism, diverse electron transport chains, and fermentation pathways is imperative for predicting accurate energy biosynthesis in Prokaryotic metabolic models. We have made core model reconstruction an essential component in our genome-scale model reconstruction pipeline by expanding previous work to reconstruct central carbon metabolism and energy biosynthesis models accurately (23). Our improved core template, which represents central carbon metabolism and energy biosynthesis, has been expanded to capture the sulfate reduction metabolism. Our previous core model template comprised the glucose oxidation pathways (Entner–Doudoroff pathway, glycolysis, pentose phosphate pathway, and the TCA cycle), aerobic and various anaerobic bacterial ETCs, and fermentation pathways (producing end products: lactate, acetate, formate, ethanol, 2,3-butanediol, butyrate, butanol, and acetone). This study expanded the core template to capture sulfate-reducing energy biosynthesis in Bacteria, hydrogenotrophic and acetoclastic methanogenesis, and methylotrophy from methanol and methylamines in Archaea.

### Quantitative prediction of ATP yield and energy biosynthesis pathways

Energy production yields are highly variable within microbial systems, depending on both growth conditions and the metabolic capabilities of each microbe. Depending on the presence or absence of various ETCs, two different microbes can display very different yields even within the same environment. Further, because the energy term in the biomass objective function of metabolic models is the most significant term by more than an order of magnitude, energy biosynthesis has an outsized effect on overall quantitative predictions from models. Given the importance of energy production for life, this system disproportionately defines the conditions in which an organism will grow and survive. For these reasons, the accuracy of a metabolic model is critically contingent on correct energy pathways. Unfortunately, these pathways can be wildly inaccurate in automatic model reconstruction pipelines and many published models; hence, we invested significant effort to improve the accuracy of the energy biosynthesis pathways within automated models constructed in MS2.

The first step to ensure that the core models accurately and quantitatively represent energy biosynthesis is to verify that the reconstructed core energy biosynthesis pathways are correctly annotated, correctly represented biochemically, and cautiously gap-filled. We developed central carbon template models comprising energy biosynthesis pathways, expanding on our previous reconstruction of central carbon metabolism across diverse microbial genomes (details in the preceding section). Our pipeline applies these templates to build a core model of each microbe of interest and tests the core model against 54 minimal media formulations, representing a broad range of energy biosynthesis life strategies. All media conditions used in this step are available in *Supplementary Table 1*. Later complications from genome-scale networks are avoided by focusing initially on the metabolic core. Production of ATP from ADP and phosphate in at least one media is verified for the reconstructed core model, where core models without such production are gap-filled to produce ATP. We select all nutrient media formulations that permit ATP production with the fewest gap-filled reactions as potential energy sources. No gap-filling is performed if the core model can produce ATP in at least one media condition. By default, at this step the core model will be gap-filled in all media formulations that require the fewest gap-filling reactions. For example, if three media conditions require one reaction to be gap-filled for ATP production during testing and all other conditions require more than two reactions to be gap-filled, the core models are gap-filled only in the three media that require one additional reaction for ATP production. Next, the core models are expanded to the genome-scale without disrupting quantitatively accurate ATP production predictions, which is achieved by testing the models in all selected energy biosynthesis nutrient formulations after adding each new reaction. If the addition of a reaction increases ATP production by over 20% above the pre-established yield for the core model in a given ATP media formulation, that reaction is rejected from being included in the model. This 20% threshold was carefully selected by repeatedly building models for over 5,000 diverse microbes with varying thresholds and reviewing the resulting ATP yield predictions.

We apply a similar approach to ensure that any gap-filling performed on our genome-scale models also does not disrupt or degrade their ATP yield predictions. We do this by prefiltering the template-specific reaction databases used to gap-fill our models in any user-specified media condition. The entire gap-filling database is added to the model simultaneously, and then any reactions that cause ATP yield to increase over 20% beyond the yield predicted by the original core model are rejected. Once filtered, the gap-filling database can then be readily and safely applied to enable the growth of the model in as many conditions as desired without having any detrimental effect on ATP predictions. Note that this process may rarely result in the gap-filling being unable to identify a working solution to permit the model to grow without breaking ATP predictions. In this case, gap-filling fails, and the user will need to manually identify an alternative solution.

Through the aforementioned procedures, the MS2 pipeline ensures that (i) all our models are constructed around a core of quantitatively accurate energy production pathways; (ii) core models can be expanded to the genome-scale without disrupting these accurate energy production predictions; and (iii) genome-scale models may be subsequently gap-filled and tested for auxotrophy without disrupting ATP predictions.

## Results and Discussion

### ModelSEED 2 genome-scale metabolic model reconstruction pipeline

The improved MS2 pipeline for genome-scale metabolic models combines and modularizes all aspects of our model reconstruction approach (see Methods and Figure 1). First, the ML classifiers select from our three defined templates based on a RAST-annotated genome. After the template is selected, ATP production is tested to ensure that infinite loops are not compromising model accuracy. Next, the reconstructed model is simulated through FBA on 54 minimal media formulations, including aerobic and anaerobic conditions for several carbon sources, and various nitrate, dimethyl sulfate, and sulfate base media formulations (see *Supplementary Table 1* for the full list of media conditions tested). If no gap-filling is required for ATP production, meaning ATP is produced in at least one of the conditions tested, we expand the model to the genome scale. If gap-filling is necessary for ATP production, we gap-fill the core model in all ATP biosynthesis media formulations. Based on the results of this analysis, we determine which ATP formulations require the fewest gap-filling reactions and accept those formulations as the most probable ATP formulations for the model. We then expand the core metabolism models to the genome scale. This expansion ensures that the model does not include any reactions that raise model ATP production by more than 20% relative to the pre-established core ATP production for any accepted ATP biosynthesis condition. The GEM is then gap-filled while again ensuring that gap-filled reactions do not increase ATP production by more than 20% relative to the pre-established core ATP production. When no medium is selected, by default, the model is gap-filled in a rich media formulation (see *Supplementary Table 2*) containing all amino acids and vitamins. We name this formulation “auxotrophy media” since it contains all essential biomass precursors for which organisms are commonly auxotrophic to ensure that pathways for these crucial compounds will not be gap-filled improperly. Optionally, users can provide a custom growth media formulation that better represents the environment and known growth conditions of the organism of interest. An App is available in KBase to run the entire MS2 pipeline.

**Figure 1.**
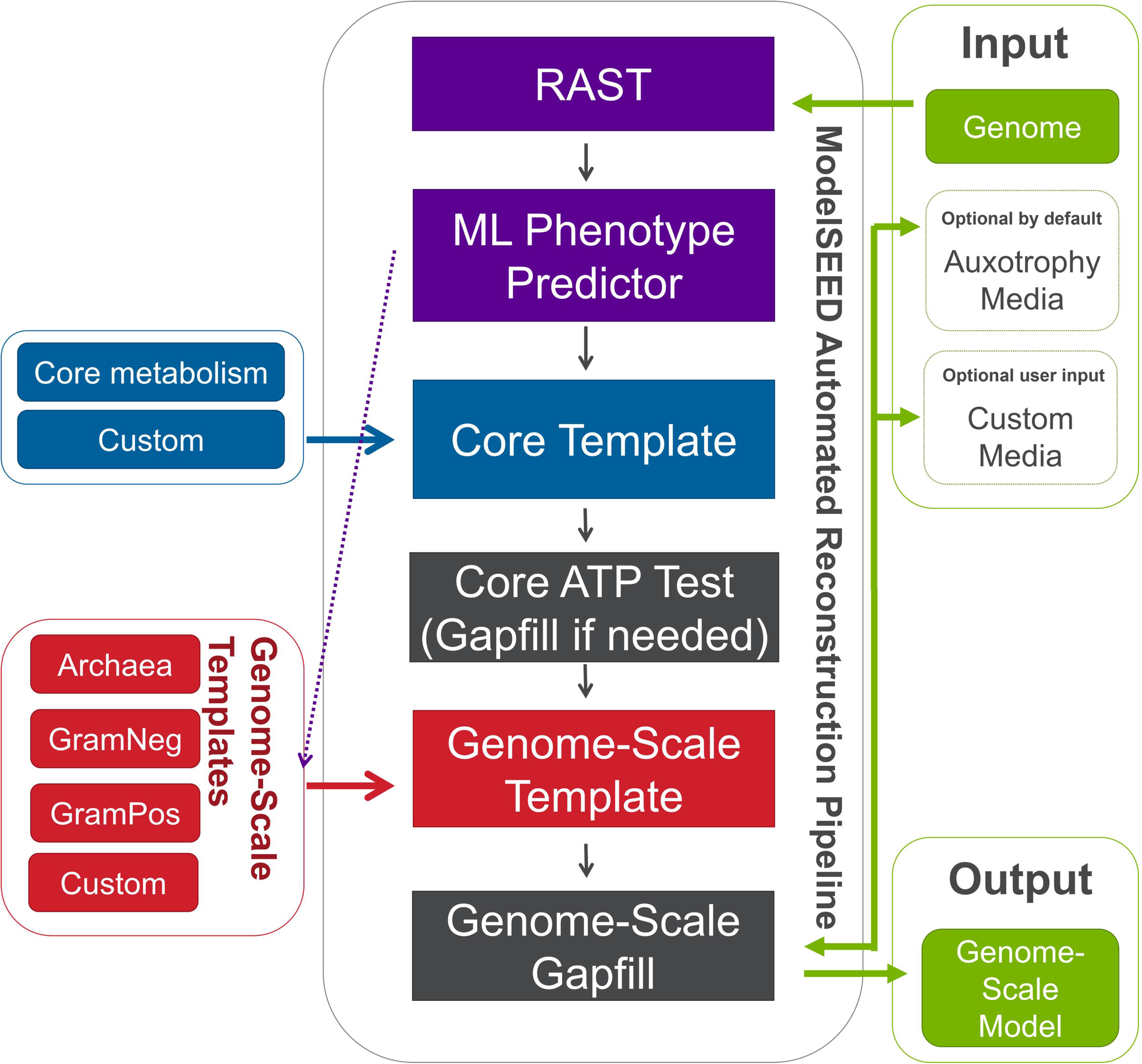
ModelSEED 2 genome-scale metabolic reconstruction pipeline enabling quantitative prediction of ATP production. A genome annotated with RAST is provided as input. Users can select a template for reconstruction, or machine-learning classifiers can select a template automatically. ATP production is tested across 54 media formulations representing diverse energy biosynthesis strategies, with gap-filling performed as necessary to ensure ATP production in at least one condition. The core metabolism model is then expanded to the genome scale, followed by gap-filling in the default auxotrophy medium or an optional user-specified medium.

We applied our new MS2 pipeline to build and analyze models for Bacteria and Archaea organisms available in KEGG (using RefSeq versions of these genomes available in KBase), for a total of 5,540 genomes, to assess improvements to the ModelSEED reconstruction pipeline. For the genome-scale study we excluded cyanobacteria, reducing our set in that analysis to 5,420 organisms. These genomes were removed from the genome-scale study because of the lack of a cyanobacteria-specific template. The analysis of cyanobacteria in the core model study in Figure 2 supports the removal of these genomes from the genome-scale study, with cyanobacteria models requiring large amounts of gap-filling at the core level; this is indicative that the core template lacks the proper pathways to represent the energy biosynthesis strategies of these organisms accurately. All genome-scale models were gap-filled with the default auxotrophy media and glucose minimal media (GMM). In addition, we built models with the previous ModelSEED template and pipeline in GMM for comparison with MS2. All GEMs in this study are available in KBase narratives listed in *ModelSEED 2 Manuscript Data* KBase Organization (https://narrative.kbase.us/#/orgs/ms2-manuscript), and all core metabolism models are available in *Supplementary Data S2*.

**Figure 2.**
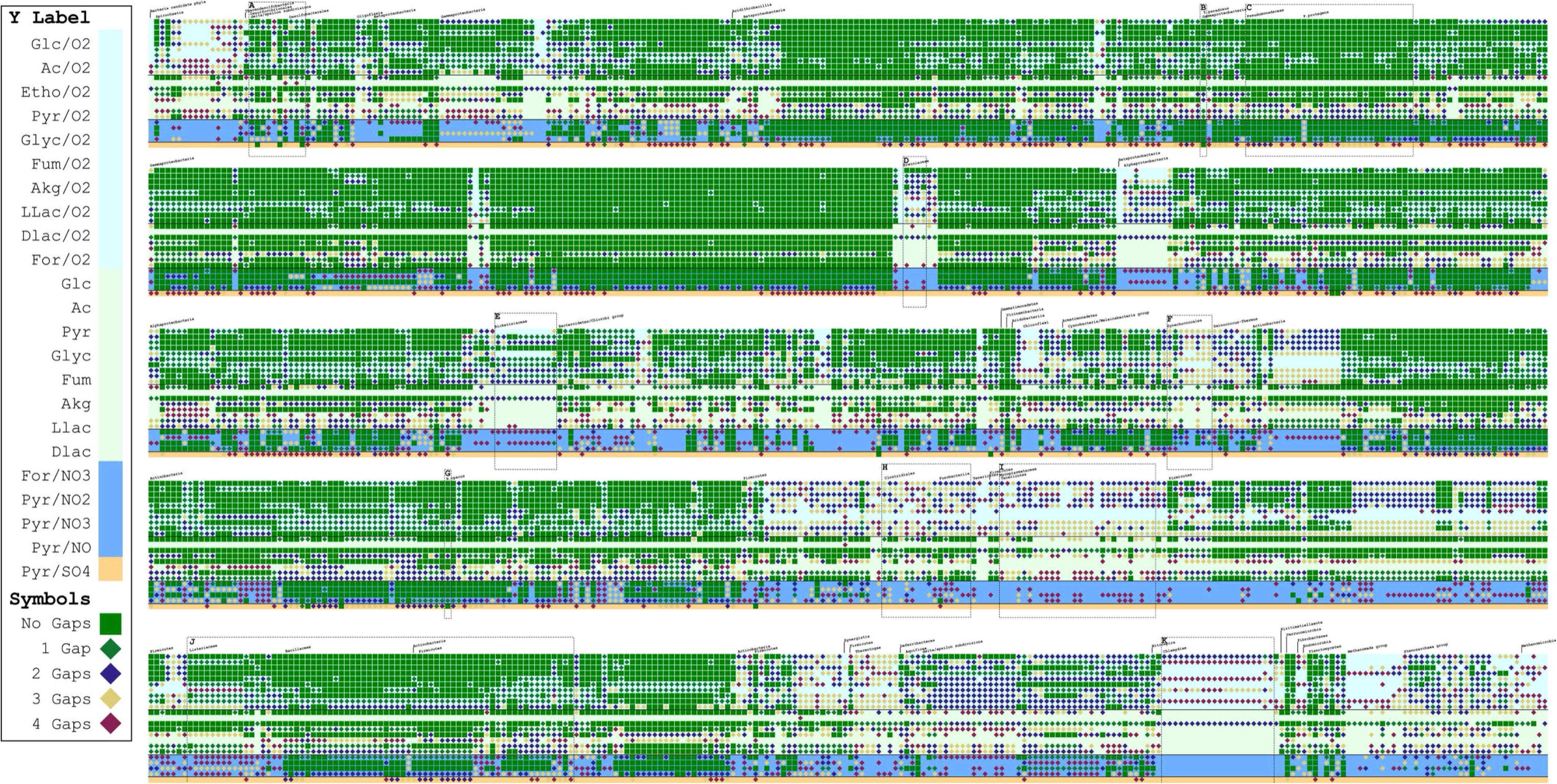
Gap-filling analysis for energy biosynthesis across diverse genomes. Green squares indicate that gap-filling is not required for ATP production in a given medium. Diamonds indicate that gap-filling is required: green, one reaction; dark blue, two reactions; yellow, three reactions; and dark pink, four reactions. The absence of a square or diamond indicates that five or more reactions are required. Light blue and light green backgrounds indicate oxic and anoxic conditions, respectively; dark blue indicates anoxic nitrate media and orange indicates anoxic sulfate media. Dashed boxes identify groups discussed in the text: A, Desulfovibrionales and Desulfobacterales; B, *Thioalkalivibrio paradoxus*; C, Pseudomonadaceae; D, Erwiniaceae; E, Rickettsiaceae; F, Synechococcales; G, *Rhodococcus opacus*; H, *Clostridium* and *Fusobacterium*; I, Mycoplasmataceae; J, Bacillales; and K, Chlamydiales. The figure shows 1,250 bacterial and archaeal genomes from the KEGG genome set. Abbreviations: Glc, glucose; Ac, acetate; Eth, ethanol; Pyr, pyruvate; Glyc, glycine; Fum, fumarate; Akg, alpha-ketoglutarate; LLac, L-lactate; DLac, D-lactate; For, formate.

### Analysis of MS2 models to study variation in energy biosynthesis pathways across diverse phylogeny

We analyzed energy biosynthesis in the core models generated as an intermediate step in the MS2 reconstruction process. We determined how much gap-filling (if any) is necessary for ATP production in the core models for our set of diverse genomes for each of the 54 media conditions tested. The full results are available in a heat map containing all 5,540 models in *Supplementary Figure 1*, from which a subset of 1,250 randomly selected genomes in 23 (of the 54) conditions tested is depicted in Figure 2. The entire ATP analysis report for all 54 media conditions is provided as output in the new MS2 App, with detailed ATP production values, gap-filled reactions, and reactions filtered out of the reconstruction to prevent ATP overproduction by the model. An example of this report is available in the MS2 Tutorial in *Supplementary KBase Narrative 1*.

Figure 2 shows the diversity of energy biosynthesis strategies across phylogeny captured by MS2 core metabolism models. To provide an overview at this scale, we reduced the level of detail in Figure 2. A large-scale version of Figure 2 is available in *Supplementary Figure 2* to facilitate more detailed analysis. Figure 2 shows that most of the core metabolism models can produce ATP in multiple media formulations without requiring any gap-filling (green squares in Figure 2). More precisely, 91% of the core models do not require any gap-filling for ATP production in at least one of the media formulations tested. Furthermore, 8% of the models have at least one medium requiring only one gap-filled reaction to produce ATP (green diamonds in Figure 2), and the remaining 1% require two gap-filled reactions (dark blue diamonds in Figure 2). These results bestow confidence that the current core metabolism template and the associated genome annotations by RAST are accurate. Gap-filling at the core metabolism step will occur only for a few complete genomes.

For some organisms, two or more gap-filling reactions are always necessary for the core metabolism model to produce ATP in aerobic media (these media formulations have a light blue background in Figure 2), which suggests that these organisms are obligate anaerobes. The best (least gap-filled) solution for most of these organisms includes fermentation pathways, including pyruvate fermentation, which further supports that most of these organisms are facultative anaerobes. For example, two of the phylogenetic groups shown in Figure 2 that require two or more gap-fill reactions for most anaerobic media formulations are obligate anaerobes: Clostridium and Fusobacterium (Box H in Figure 2) (36, 37).

Nitrate-reducing bacteria are naturally abundant and phylogenetically diverse, with over 50 genera (38). This diversity is evident in Figure 2, where a spectrum of ATP production is observed in nitrate-containing anaerobic media sources (dark blue background) without any gap-filling requirements. In the Bacillales order, we observe that many Bacillus species with nitrogen fixation capabilities (39, 40) (Box J in Figure 2) are predicted to produce ATP in the nitrate media formulations, while many Listeria species are pathogens without nitrate-reducing capabilities reported in the literature (left of the Bacillus species in Box J in Figure 2) and require gap-filling to produce ATP in nitrate media formulations. The Pseudomonadaceae family (Box C in Figure 2) shows that every Pseudomonas species except *Pseudomonas protegens* – a strict aerobe that is known not to reduce nitrate (41) *–* can reduce nitrate media sources without gaps. These results suggest that MS2 correctly predicts nitrate reduction when building models with our latest pipeline.

With the introduction of sulfur reduction energy biosynthesis and respective ETCs, we expect to find organisms across a diverse phylogenetic range capable of reducing sulfate without any gap-filling. Members of the Desulfovibrionales and Desulfobacterales orders in the first row of Figure 2 (Box A) depict three organisms that require no gap-filling for growth in “Pyr/SO4” (sulfate minimal) media, which is expected since the majority of members of these orders are sulfur-reducing bacteria (42). Other organisms in these taxonomic groups require gap-filling to grow in sulfate minimal media. This may be attributed to a lack of other sulfur sources in testing, such as sulfite or thiosulfate, variations in sulfur-reduction metabolism not captured in our core template, or poor annotation of sulfur-reducing genes. The last possibility seems likely when we observe no issues with well-studied species in these taxonomic groups, such as *Desulfovibrio vulgaris str. Hildenborough* (43). Sulfate reduction is additionally correctly predicted in phylogenetically diverse organisms such as *Thioalkalivibrio paradoxus* (44) (Gammaproteobacteria, Box B in Figure 2) and *Rhodococcus opacus* (45) (Actinomycetia, Box G in Figure 2).

The aforementioned analysis can also identify phylogenetic groups where pathway representation or energy biosynthesis strategies can be further improved (e.g., the areas of Figure 2 that require extensive amounts of gap-filling across most media formulations). An example of such cases is the Synechococcales order (Box F in Figure 2), where five or more reactions must be gap-filled for ATP production. This result identifies cyanobacteria as a group for which the current templates are not sufficient and a specialized template would be required. The high core-level gap-filling requirement serves as a diagnostic warning that the current template does not adequately represent cyanobacterial energy metabolism. Users should therefore review and curate cyanobacterial models before applying them in downstream studies. We observe similar high gap-filling requirements for members of the Mycoplasmataceae family (Box I in Figure 2), whose unique physiology may also require a specialized template for accurate energy biosynthesis representation. The modular MS2 pipeline permits specialized templates to be substituted into the existing framework without requiring refactoring of the pipeline.

Extensive gap-filling is acceptable in some cases, such as the Buchnera genus members of the Erwiniaceae family in Figure 2 Box D, which are endosymbionts living inside aphid guts (46). In such an environment, where the critical processes for survival are shared with the host, a community model of symbiont and host interactions is necessary to model these organisms adequately, but extensive gap-filling is needed to simulate the isolated endosymbiont. Other examples include members of the genus Wolbachia (Rickettsiaceae family in Figure 2 Box E), which are intracellular parasites that infect arthropods and filarial nematodes (47), and Chlamydiales, which are obligate intracellular bacteria that live in insects, protozoa, and animals such as the human pathogen *Chlamydia trachomatis* (48). These organisms share the same gap-filling profile in Box K in Figure 2.

The acetate core media formulation was an important negative control in template testing where, without an electron acceptor, no organism is expected to produce ATP in this media. The expected negative control result is depicted in Figure 2 as the “Ac” (acetate) media condition, which is predictably a blank line except for those genomes that require at least four gap-filled reactions.

### Comparison of MS2 models with CarveMe models

A qualitative and quantitative assessment of the ATP yield predictions by the MS2 pipeline was conducted, focusing only on those models where ATP production was predicted with no core gap-filling required. This approach reveals how the MS2 core template represents energy biosynthesis without gap-filling support. These predictions were compared with those generated from models built with the CarveMe pipeline (*Supplementary Data S3*) by building models of the same set of diverse genomes through CarveMe and MS2.

All gap-filling reactions from the CarveMe models were removed to directly compare with the MS2 templates without the interference of gap-filling contributions. This comparison evaluates the core models since testing of energy biosynthesis in the MS2 pipeline occurs before gap-filling. ATP production of MS2 core models does not significantly differ when expanded to genome-scale since the MS2 prevents genome-scale ATP production from exceeding >20% of the core model ATP production. We did not use CarveMe’s specialized cyanobacteria template for this comparison since the ModelSEED still lacks a comparable template; hence, CarveMe is better suited for reconstructing cyanobacteria models, particularly if users know that an organism is within this taxonomic group before reconstruction. Pre-reconstruction identification of organisms can be challenging, particularly in modeling numerous genomes or MAGs, and requires a separate set of specialized tools. Figure 3 illustrates this comparison for some key phylogenetic groups, while the complete comparison for 5,420 genomes is available in *Supplementary Figure 3* and *Supplementary Figure 4*.

**Figure 3.**
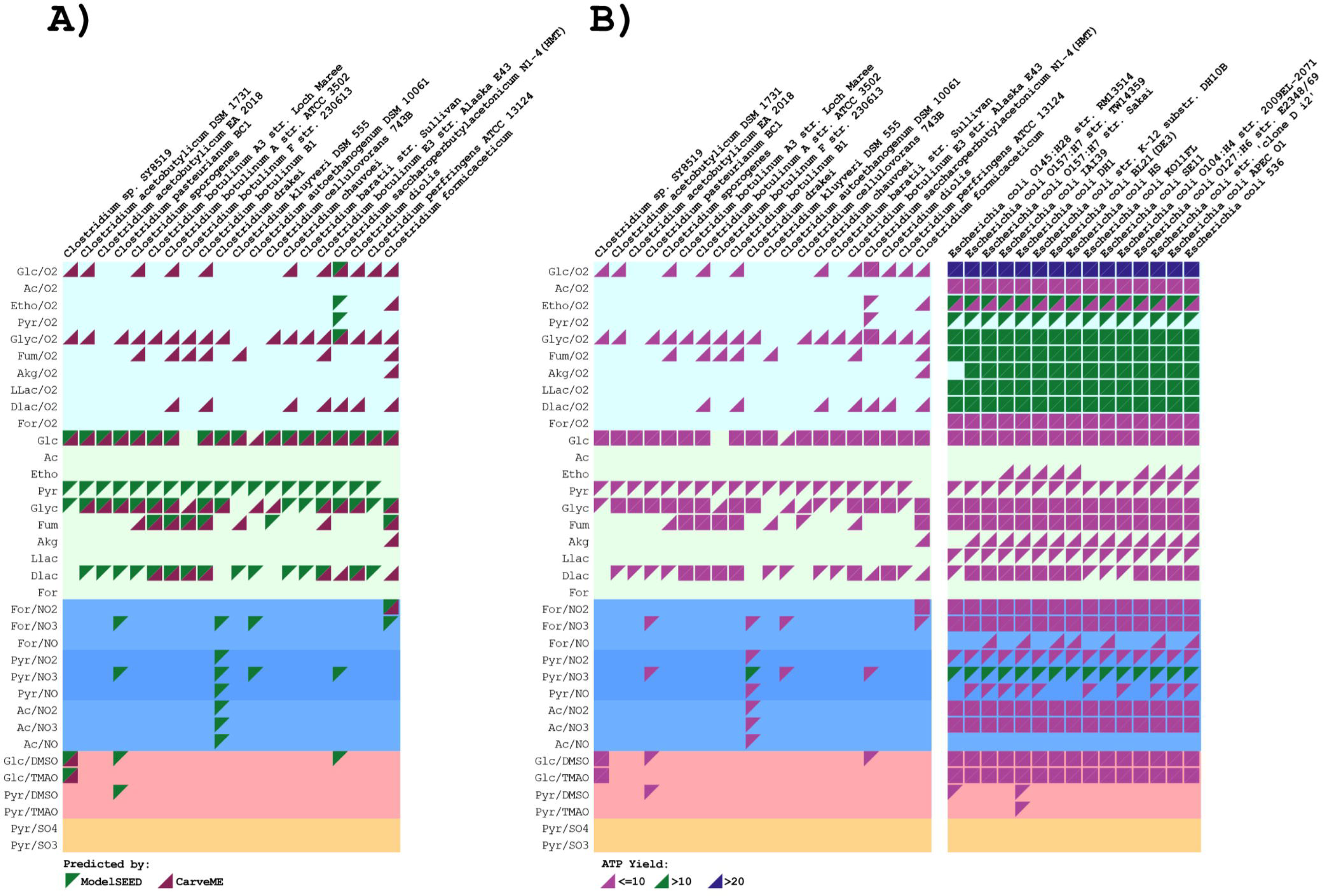
Qualitative and quantitative comparison of ATP predictions from ModelSEED and CarveMe models. (A) Qualitative ATP predictions for 20 members of the Clostridiaceae family. A triangle indicates that a model produces ATP in the specified medium; green indicates ModelSEED and purple indicates CarveMe. (B) Quantitative comparison for 50 Gammaproteobacteria. The upper triangle represents ModelSEED and the lower triangle represents CarveMe. Colors indicate predicted ATP yield: pink, 0-10; green, 10-20; and blue, greater than 20 mol ATP per mol nutrient. Abbreviations are as in Figure 2.

The predictions from MS2 and CarveMe generally exhibit high similarity across phyla. A statistical assessment of this agreement for various taxa is available in *Supplementary Tables 3A, B, and C*. In Figure 3A, ATP prediction results for a random subset of 20 members of the Clostridiaceae family are compared. The presence of a triangle in Figure 3A signifies that a model can produce ATP (any amount of ATP allowed) for a given media formulation. The upper triangle in Figure 3A represents the ModelSEED predictions in green, and the lower triangle represents CarveMe predictions in purple. The absence of triangles indicates that neither MS2 nor CarveMe predicts any ATP biosynthesis for that organism and media condition. We observe a good level of agreement between the two pipelines where ATP production is not predicted for many media conditions across genomes by either MS2 or CarveMe. ATP production predictions in anaerobic conditions (light green background) also show a marked concordance, particularly in glucose (Glc), glycogen (Glyc), and fumarate (Fum) minimal media. This is expected because members of the genus Clostridium are primarily obligate anaerobes (36). CarveMe, on the other hand, also makes predictions of ATP production in aerobic media formulations (light blue background), whereas MS2 predicts no ATP production in these conditions, as expected for obligate anaerobes. The exception here is the butanol-producing bacterium *Clostridium saccharoperbutylacetonicum N1-4* (an obligate anaerobe (49)), for which both MS2 and CarveMe predict aerobic ATP biosynthesis. Upon further analysis, for all 63 members of the anaerobic Clostridiaceae family (see *Supplementary Figure 3*), MS2 predicts ATP production only in aerobic media for C. *saccharoperbutylacetonicum N1-4.* A detailed comparison of the annotations with other family members is necessary to identify the root cause of this discrepancy. This organism can also be a case study to test and tune our methods for predicting anaerobic vs. aerobic growth (see methods section) and adjust thresholds accordingly. CarveMe models for other known obligate anaerobes, such as Fusobacterium, can also utilize oxygen to produce ATP (see *Supplementary Figure 3*).

Figure 3B shows the same subset of Clostridium genomes with plotted data for quantitative analysis. The colors represent different ranges of predicted ATP yields: pink, 0 to 10; green, 10 to 20; blue > 20. Clostridium’s ATP production within the lower range of 0 to 10 for all media formulations is consistent with these organisms’ energy biosynthesis as obligate anaerobes. In contrast, a random subset of members of the Escherichia genus in Figure 3B exhibit the greatest ATP production of >20 mol ATP per mol nutrient consumed for aerobic glucose media, versus between 10 and 20 for most other aerobic media formulations. Members of the Escherichia genus are among the most studied microbes, with high-quality annotations begetting high-quality metabolic models; therefore, unsurprisingly, both MS2 and CarveMe exhibit agreement across most media formulations with these organisms. There is a discrepancy, however, with ATP production in aerobic conditions across the entire dataset shown in Figure 4. MS2 ATP production in glucose (Glc/O2) media has a median value of ∼26 ATP, with most of the predicted values showing little variation for the 25 to 75 percentiles, while CarveMe exhibits more variation and a median value below 20.

**Figure 4.**
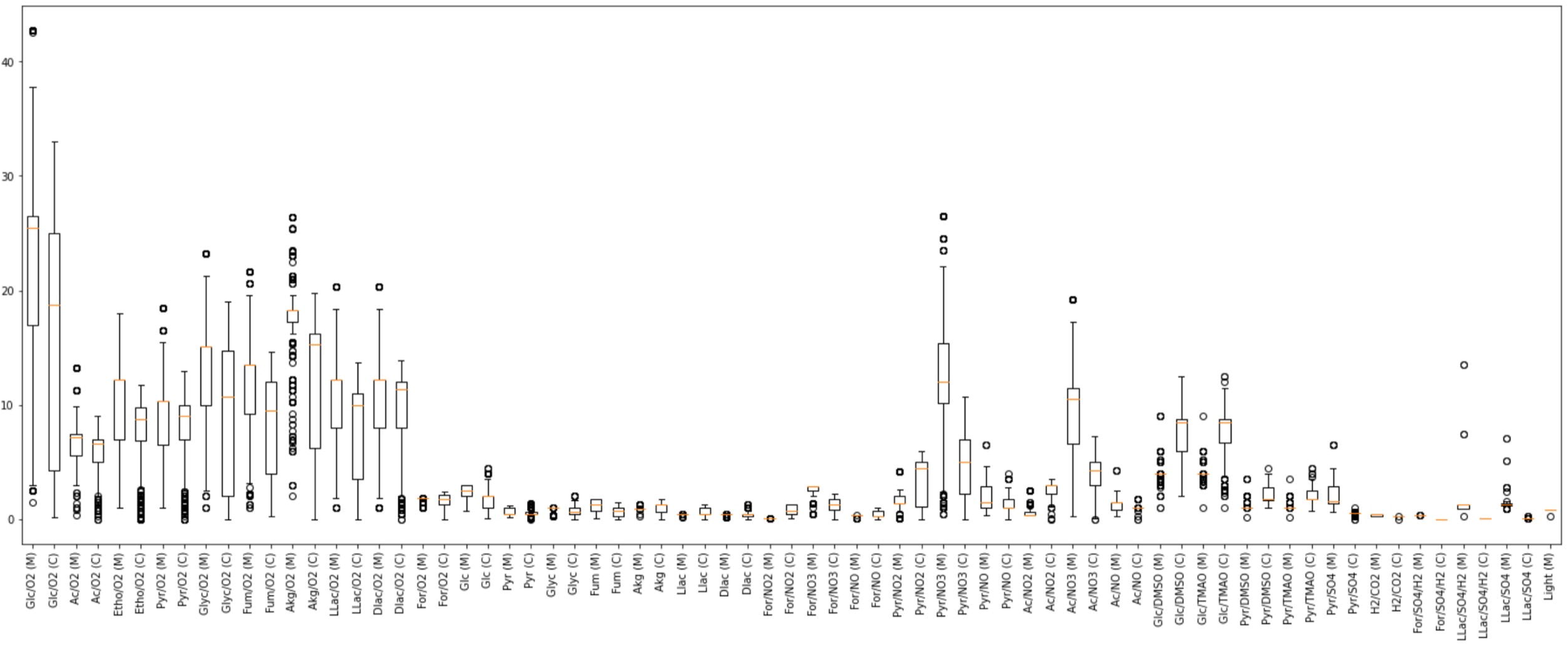
ATP yield distributions for each medium across ModelSEED and CarveMe models. Boxplots show ATP yield per unit of carbon source for models generated by ModelSEED (M) and CarveMe (C). Abbreviations: Glc, glucose; Ac, acetate; Etho, ethanol; Pyr, pyruvate; Glyc, glycerol; Fum, fumarate; Akg, alpha-ketoglutarate; LLac/DLac, L-/D-lactate; For, formate.

We observe the same behavior for other carbon sources in aerobic media formulations where MS2 predicts a higher median value of ATP with less variation for the 25 to 75 percentiles than CarveMe. This is most notably observed in glycerol (Gly/O2), fumarate (Fum/02), α-ketoglutarate (Akg/O2), and D-lactate (Dlac/O2) media.

Figure 3A also illustrates ATP production in nitrogen minimal media for a handful of organisms, mainly by MS2; however, there does not appear to be evidence of nitrate reduction for these organisms (including the previously described C. *saccharoperbutylacetonicum N1-4* “problematic” genome). MS2 and CarveMe mostly agree for the Pseudomonadaceae with known nitrate-reducing members, although MS2 predicts ATP production without gap-filling for more members than CarveMe, and MS2 predicts ATP production in at least one nitrate base media formulation for 127/136 genomes in the Bacillus genus versus 10/136 from CarveMe. For the entire Bacillales order, MS2 predicts growth in at least one nitrate reduction media in 324 more genomes than CarveMe (*Supplementary Table 3C*). Figure 4 also shows the ATP production range for nitrate media formulations. ModelSEED furthermore predicts growth in nitrate media (Pyr/NO3 and Ac/NO3) for more organisms identified as nitrate reducers than CarveMe.

### Improvements in MS2 models compared with the previous ModelSEED

We reconstructed the same set of 5,420 genome-scale models with the previous version of the pipeline and legacy templates for the same set of RAST annotated genomes. We note that legacy templates were optimized for older versions of the RAST annotations pipelines. The changes presented in Figure 5 reflect not only the evolution of our templates but also the evolution of the annotations in the RAST pipeline over time.

**Figure 5.**
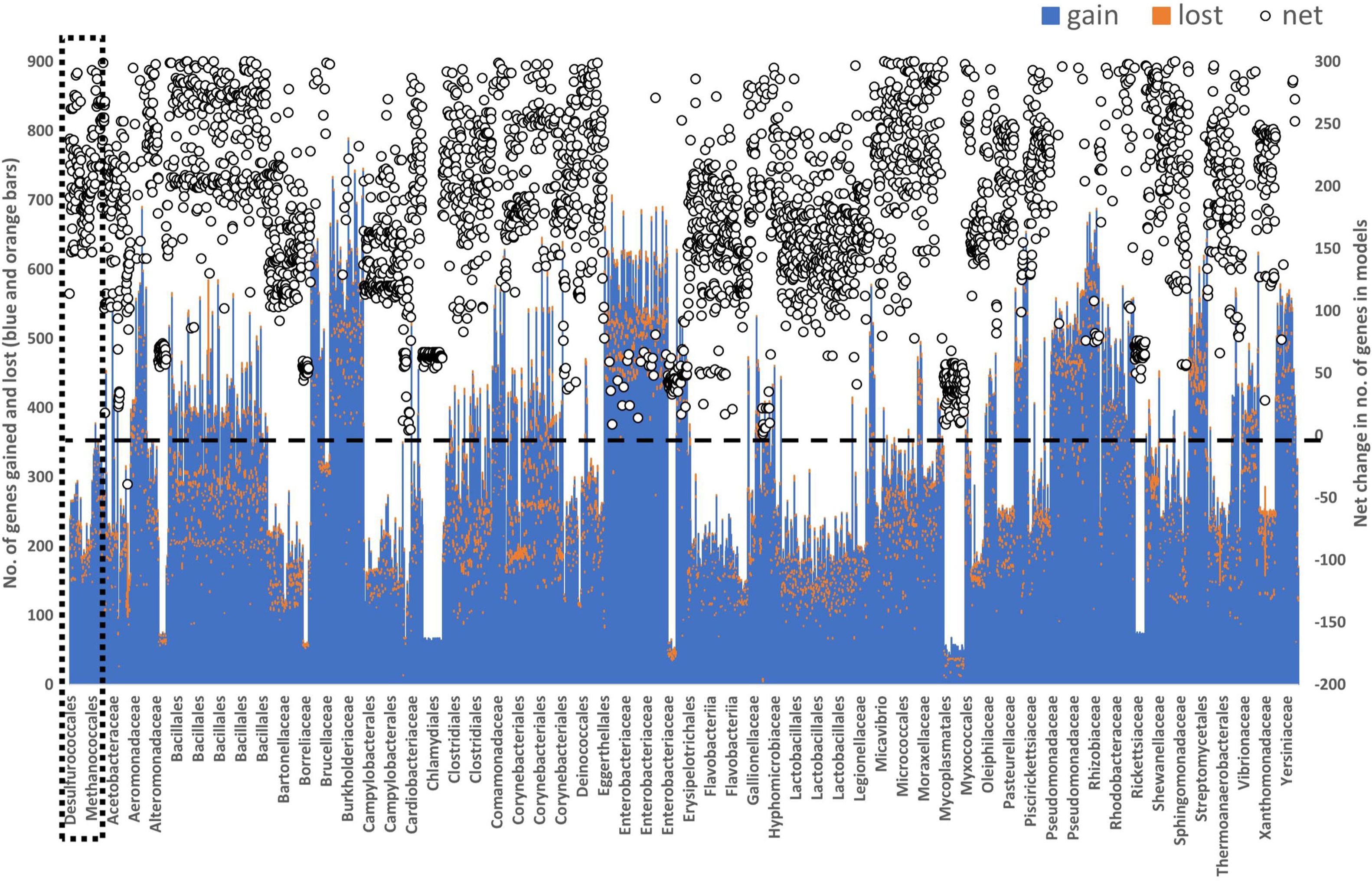
Comparison of the legacy ModelSEED pipeline and templates with the ModelSEED v2 pipeline and templates. On the primary vertical axis, stacked bars show genes gained in ModelSEED v2 models (blue) and genes lost relative to legacy ModelSEED models (orange). Circles on the secondary vertical axis show the net change in model-associated gene count for each organism. The dashed box highlights archaeal genomes.

The number of genes included in both pipeline versions was compared to assess the impact of the introduced changes without gap-filling. Both sets of models were gap-filled in GMM, although, gap-filled reactions are not mapped to genes. Genes were generally gained in our models across all phylogenetic groups. A significant increase in Archaeal genes (dashed-line box in Figure 5) is observed because MS2 expanded Archaeal annotations and introduced an Archaea-specific template. Poorly mapped annotations explain many circumstances when genes were lost, where knowledge of gene-protein-reaction relationships needs to be updated. Transporter annotations are especially challenging; hence, hundreds of gene functional role annotations associated with transport reactions were manually mapped and refined for the MS2 pipeline. This improvement introduced more specific annotations to the detriment of older, more generic, annotations such as those where EC numbers were outdated. Lumped reactions were also divided into multiple steps, each defined by a gene-to-SEED functional role mapping. The latest version of the ModelSEED Biochemistry database, where mass-balanced reactions have almost doubled since the original 2010 release (28), was also applied for these efforts. Significant emphasis was placed on the manual curation of every reaction in our core metabolism templates, which substantially improved upon the previous energy biosynthesis analysis. Comparing ATP production profiles for both versions, MS2 importantly eliminates infinite ATP production caused by energy loops. The full dataset used to plot Figure 5 is available in *Supplementary Table 4*.

### MS2 model comparison in glucose minimal and auxotrophy media

Beyond studying ATP biosynthesis strategies, the models produced by our MS2 pipeline also reveal insights into the likelihood of auxotrophic dependencies among our selected collection of 5,420 diverse genomes. We gained these insights by comparing the models generated by MS2 across two gap-filling conditions: GMM and auxotrophy media. In both cases, we start with the same genome and annotations, meaning the core metabolism, ATP biosynthesis, and gene-associated reactions will be identical in both cases. However, we see that models gap-filled in GMM consistently have more gap-filled reactions than models gap-filled in auxotrophy media (Figure 6). The reason is that auxotrophy media provides most essential biomass components in the environment and, therefore, models require less gap-filling to complete intracellular biosynthesis pathways for these same compounds. In contrast, GMM provides only minimal nutrients, meaning all biosynthesis pathways that produce essential metabolites are forced to be gap-filled. The difference in the number of gap-filling reactions between the GMM and auxotrophy models tells us how far each model is, based on annotated gene-associated reactions only, from being able to produce all essential metabolites. There are three interpretations for these differences: (1) a large difference indicates the organism associated with the model has more auxotrophic dependencies; (2) the genome associated with the model is poorly annotated and simply has more gaps in all of its pathways; or (3) the genome in question has a distinctive metabolism that is not well captured by the template we used to build the model. Fortunately, we can differentiate scenario (2) and (3) from scenario (1) based on how many reactions were gap-filled in the model overall (e.g., how many reactions were gap-filled in auxotrophic media or in the core). For this reason, we included the difference between the GMM and auxotrophy gap-filling counts normalized by the GMM gap-filling counts as a third data element in Figure 6 (red points, second axis). If this normalized gap-filling difference is close to 1, then the organism is more likely to be highly auxotrophic; if the number is close to zero, then the organism is likely to grow in near-minimal media. This analysis reveals high normalized differences in taxonomic groups composed of endosymbionts or parasitic intracellular organisms (e.g., Rickettsiales, Mollicutes, Chlamydia), where auxotrophy is common since these species rely extensively on host metabolism, validating this approach. We note, this rough approach lacks the resolution needed to predict specific auxotrophic dependencies, but it does provide a meaningful measure of the overall level of auxotrophy that an organism might have.

**Figure 6.**
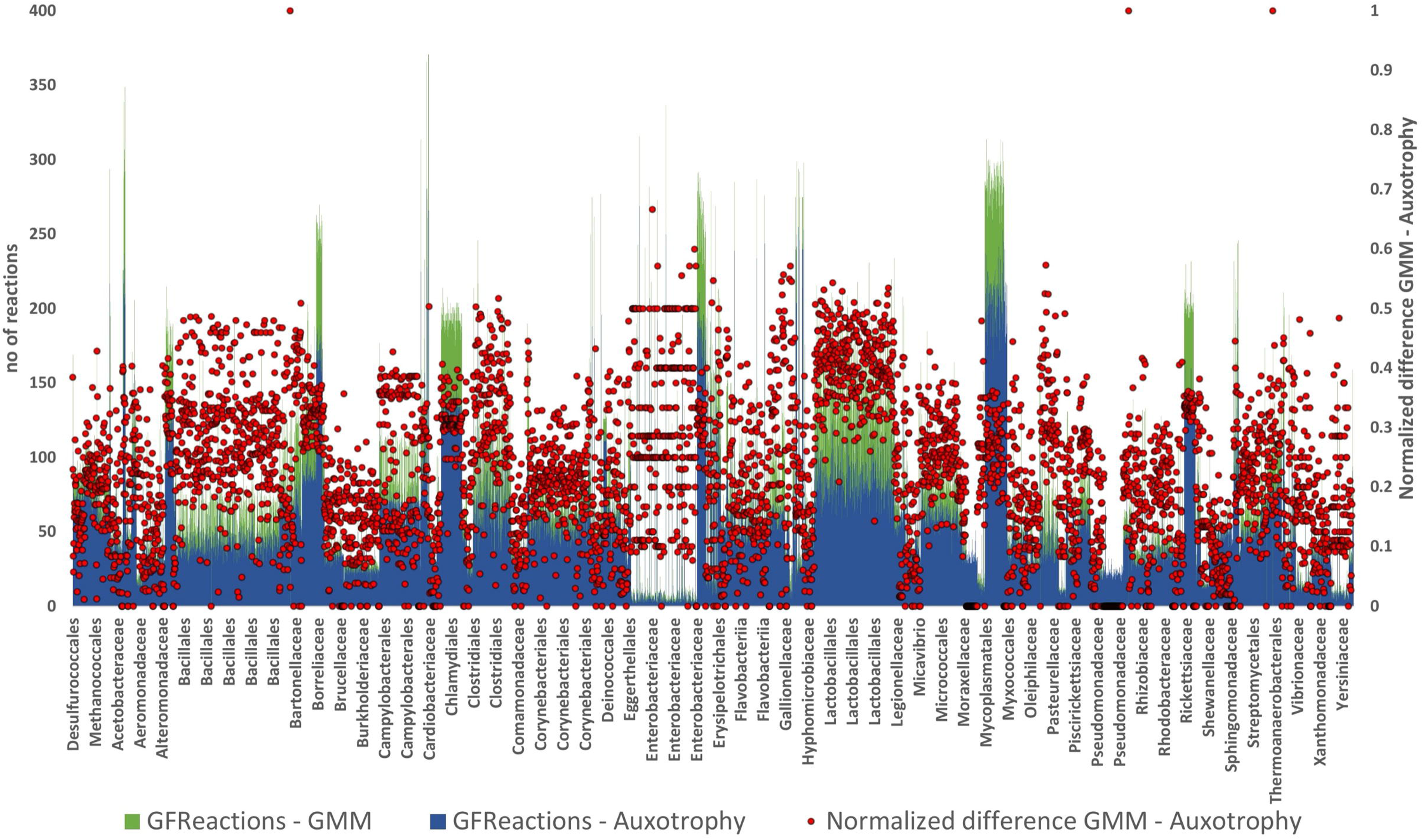
Comparison of models gap-filled in glucose minimal medium and auxotrophy medium. Total gap-filled reactions are shown for models of 5,420 genomes. Models gap-filled in glucose minimal medium (GMM) are shown in green and models gap-filled in auxotrophy medium are shown in dark blue. Red points on the secondary axis show the difference between GMM and auxotrophy gap-filling counts normalized by the GMM gap-filling count.

For the genomes that fall into the poorly annotated category by having large numbers of gap-filled reactions overall, we were able to use this analysis in our curation efforts to inspect the most common gap-filled reactions and to curate their gene role-to-reaction mappings when appropriate. Based on this analysis, we were able to identify and correct many consistent annotation or template errors, reducing the needed gap-filling between our early templates and our final templates by 57% on average.

We note that gap-filling is not always a sign of poorly annotated and thus incomplete pathways. High gap-filling can also indicate errors in the biomass objective function of the model or errors in our classification of the genomes, resulting in the wrong template (and thus the wrong biomass reaction) being used to build the models. The most prominent example of this in Figure 6 is the mycobacteria family. Since we lack a specific template for this family at this time, these species are all currently classified as gram negative. Since these microbes completely lack any sort of cell wall, this will result in the models gap-filling the entire gram-negative cell wall biosynthesis pathway. We can see that the large number of reactions in the species validates this prediction. In the future, we can associate specific biomass compounds with each gap-filled reaction to determine whether specific biomass compounds are to blame for most gap-filling in certain clades, and this data can then be used to systematically refine the biomass reactions for all our models. The full dataset used to plot Figure 6 is available in *Supplementary Table 5*.

## Conclusions

In this work, we comprehensively updated the ModelSEED reconstruction pipeline and present the ModelSEED 2 (MS2) as the first major release since the original 2010 publication (Figure 1). Core model reconstruction is now a critical step in the MS2 reconstruction pipeline, making ATP production more accurate at the core level and preventing ATP overproduction when expanding to the genome scale and in any subsequent gap-filling. The energy biosynthesis analysis (Figure 2) revealed the predictive capability of our core templates across diverse media and ATP production in anaerobic vs. aerobic conditions. The ATP-safe method was demonstrated to accurately describe obligate anaerobes and strict and facultative aerobes. Accurate predictions of nitrate and sulfate reducers were demonstrated as well.

The MS2 pipeline was compared (Figures 3 and 4) with the CarveMe pipeline (a top-performing modeling reconstruction resource) to assess improvements in our capture of aerobic and anaerobic respiration in core models. Overall, we observe a lot of concordance between the two platforms, with MS2 showcasing its strength in accurate predictions of energy biosynthesis. We expect the template approach of MS2 to perform better across phylogeny and for incomplete genomes. CarveMe will still outperform for organisms that are phylogenetically close to those used to generate its universal BiGG model. This comparison also illuminated improvements that are still necessary, most notably in cyanobacteria reconstruction, which were echoed when assessing the observations from the core models (Figure 2). We also compared models from MS2 to those generated through the old ModelSEED pipeline, which revealed that MS2 models possess more genes and on average 57% less gap-filled reactions. The study results plotted in Figure 6 also illustrate the limitations of using universal biomass equation definitions, where endosymbiont and parasitic organisms require many gap-filled reactions to satisfy universal biomass requirements fulfilled in vivo by the host. The universal biomass reaction also represents multiple quinones due to the difficulty in annotating specific quinones across phylogeny, which materializes in gap-filling for quinone biosynthesis in organisms that may not have a specific quinone in their genome. Similar results are expected for other biomass cofactors that vary across phylogeny, are represented in the universal biomass reaction, and whose biosynthesis may be incorrectly gap-filled. Future improvements to the ModelSEED pipeline will prioritize improvements in biomass equation specificity to compensate for these shortcomings.

RAST continues to be our primary source of curated annotations to reaction mappings because of the consistency in its controlled vocabulary (50). Nevertheless, combining multiple annotation platforms may better compensate for some notable weaknesses of RAST (51). This functionality of blending annotation platforms is already available in KBase. The KBase infrastructure of the MS2 pipeline also supports third-party developers to improve the method, for example, allowing users to upload annotations from various supported sources and directly substitute them into MS2. Together with the ability to run the reconstruction pipeline on custom templates, we want to empower the community to actively contribute to areas where we fall short and in niches that suit their research needs. We believe this joint development with the community is the future for the ModelSEED reconstruction pipeline. In the MS2 redesign, we deliberately fostered the development of third-party tools by modularizing every pipeline step to incentivize community-developed add-ons. We expect the community to introduce ML classifiers, new templates, and complementary analysis tools to the pipeline. The redesigned ModelSEED website provides direct access to reconstruction, gap-filling, flux balance analysis, model editing, and model export. Its open-source backend is available from the ModelSEED API repository (https://github.com/ModelSEED/modelseed-api) and can also be deployed locally in a Docker container, supporting reproducible programmatic and web-based use of ModelSEED. The API uses ModelSEEDpy and COBRApy (52) as components of its modeling stack.

## Acknowledgements

National Science Foundation MCB-1716285

This work is supported as part of the BER Genomic Science Program. The DOE Systems Biology Knowledgebase (KBase) is funded by the U.S. Department of Energy, Office of Science, Office of Biological and Environmental Research under Award Numbers DE-AC02-05CH11231, DE-AC02-06CH11357, DE-AC05-00OR22725, and DE-AC02-98CH10886.

The submitted manuscript has been created by UChicago Argonne, LLC as Operator of Argonne National Laboratory (‘Argonne’) under Contract No. DE-AC02–06CH11357 with the U.S. Department of Energy. The U.S. Government retains for itself, and others acting on its behalf, a paid-up, nonexclusive, irrevocable, worldwide license in said article to reproduce, prepare derivative works, distribute copies to the public, and perform publicly and display publicly, by or on behalf of the Government. The Department of Energy will provide public access to these results of federally sponsored research in accordance with the DOE Public Access Plan http://energy.gov/downloads/doe-public-access-plan.

## Supplementary Data

Supplementary Tables and Figures, all core metabolic models, and CarveMe models are available at https://bioseed.mcs.anl.gov/~fliu/modelseed2/. The revised supplementary files selected during bioRxiv submission correspond to this version of the manuscript.

*ModelSEED 2 Manuscript Data* KBase Organization (https://narrative.kbase.us/#/orgs/ms2-manuscript)

*Supplementary KBase Narrative 1* - MS2 Tutorial

*Supplementary Table 1 -* Minimal media formulation for ATP production testing.

*Supplementary Table 2* - Auxotrophy media composition.

*Supplementary Table 3 A, B, and C* - CarveMe comparison summary stats.

*Supplementary Table 4* - Supporting data for Figure 5.

*Supplementary Table 5* - Supporting data for Figure 6.

*Supplementary Figure 1* - Gap-filling analysis for energy biosynthesis across diverse genomes for the complete set of 5,540 genomes.

*Supplementary Figure 2* - Gap-filling analysis for energy biosynthesis across diverse genomes for the complete set of 1,250 genomes.

*Supplementary Figure 3* - Qualitative assessment of the ATP prediction results for ModelSEED and CarveMe metabolic models for the complete set of 5,420 genomes.

*Supplementary Figure 4* - Qualitative assessment of the ATP prediction results for ModelSEED and CarveMe metabolic models for the complete set of 5,420 genomes.

*Supplementary Data S1* - Model reconstruction templates.

*Supplementary Data S2* - Core metabolism models.

*Supplementary Data S3* - CarveMe models.

Supplementary data are available with this revised bioRxiv version and at the ModelSEED supplementary-data site.

## Conflict of Interest

None declared.

## Data Availability

The ModelSEED v2 pipeline is available through the ModelSEED website (https://modelseed.org/) and KBase (https://narrative.kbase.us/#catalog/apps/ModelSEEDReconstruction/build_metabolic_models). The current ModelSEED website is backed by the open-source ModelSEED API version 0.1.0 (https://github.com/ModelSEED/modelseed-api; MIT license) and has a container image for local deployment. The API uses ModelSEEDpy for metabolic modeling. Versioned model reconstruction templates are available at https://github.com/ModelSEED/ModelSEEDTemplates; the exact templates used for this study are included in *Supplementary Data S1*. The 5,540 core models and the genome-scale models analyzed in this study are available through the *ModelSEED 2 Manuscript Data* KBase organization (https://narrative.kbase.us/#/orgs/ms2-manuscript). Core models are also provided in *Supplementary Data S2*, and the CarveMe comparison models are provided in *Supplementary Data S3*. The complete dataset, including the reconstruction templates, model collections, supplementary tables, and supplementary figures, is archived in Zenodo at https://doi.org/10.5281/zenodo.21639259.

## Notes

https://narrative.kbase.us/#/orgs/ms2-manuscript

https://doi.org/10.5281/zenodo.21639259

https://github.com/ModelSEED/modelseed-api

### Competing Interest Statement

The authors have declared no competing interest.

### Summary of Updates

This version updates the manuscript to reflect the restored and operational ModelSEED infrastructure. ModelSEED v2 is now accessible through the redesigned ModelSEED website, which supports metabolic model reconstruction, gap-filling, flux balance analysis, model editing, and model export. The website is backed by the public ModelSEED API repository and supports local container deployment. The abstract, introduction, conclusions, and data availability statements were revised to replace planned-release language with the current deployment status. The software description now identifies modelseed-api as the public backend and ModelSEEDpy as part of its modeling stack. Vibhav Setlur was added as an author for software contributions to the deployed ModelSEED frontend. The template discussion and cyanobacteria results were revised to clarify that a cyanobacteria-specific template is not currently available and that cyanobacterial reconstructions require careful review and curation. Figure legends, the conflict-of-interest statement, and the data-availability and supplementary-data descriptions were updated, including a public Zenodo archive for the study data.

